# Inference of multiple-wave population admixture by modeling decay of linkage disequilibrium with multiple exponential functions

**DOI:** 10.1101/026757

**Authors:** Ying Zhou, Kai Yuan, Yaoliang Yu, Xumin Ni, Pengtao Xie, Eric P. Xing, Shuhua Xu

**Affiliations:** Chinese Academy of Sciences (CAS) Key Laboratory of Computational Biology, Max Planck Independent Research Group on Population Genomics, CAS-MPG Partner Institute for Computational Biology, Shanghai Institutes for Biological Sciences, Chinese Academy of Sciences, Shanghai, 200031, China; Machine Learning Department, Carnegie Mellon University, Pittsburgh, PA, 15213, USA; Department of Mathematics, School of Science, Beijing Jiaotong University, Beijing 100044, China; School of Life Science and Technology, ShanghaiTec University, Shanghai 200031, China; Collaborative Innovation Center of Genetics and Development, Shanghai 200438, China

**Keywords:** Population admixture, Linkage Disequilibrium (LD), Admixture model, Multiple exponential functions, SNP

## Abstract

Admixture-introduced linkage disequilibrium (LD) has recently been introduced into the inference of the histories of complex admixtures. However, the influence of ancestral source populations on the LD pattern in admixed populations is not properly taken into consideration by currently available methods, which affects the estimation of several gene flow parameters from empirical data. We first illustrated the dynamic changes of LD in admixed populations and mathematically formulated the LD under a generalized admixture model with finite population size. We next developed a new method, MALDmef, by fitting LD with multiple exponential functions for inferring and dating multiple-wave admixtures. MALDmef takes into account the effects of source populations which substantially affect modeling LD in admixed population, which renders it capable of efficiently detecting and dating multiple-wave admixture events. The performance of MALDmef was evaluated by simulation and it was shown to be more accurate than MALDER, a state-of-the-art method that was recently developed for similar purposes, under various admixture models. We further applied MALDmef to analyzing genome-wide data from the Human Genome Diversity Project (HGDP) and the HapMap Project. Interestingly, we were able to identify more than one admixture events in several populations, which have yet to be reported. For example, two major admixture events were identified in the Xinjiang Uyghur, occurring around 27–30 generations ago and 182–195 generations ago, respectively. In an African population (MKK), three recent major admixtures occurring 13–16, 50–67, and 107–139 generations ago were detected. Our method is a considerable improvement over other current methods and further facilitates the inference of the histories of complex population admixtures.

## Introduction

The “Out of Africa” human migrations resulted in population differentiation in different continents, while subsequent migrations that have occurred over the past millennia have resulted in gene flow among previously separated human sub-populations. As a result, admixed populations come into being when previously mutually isolated populations meet and intermarry. Population admixture has received a great deal of attention recently. Many studies based on genome-wide data have shown that gene flow has been common among inter-continental and intra-continental populations and that admixture of populations often leads to extended linkage disequilibrium (LD), which can greatly facilitate the mapping of human disease genes(McKeigue 2005; Reich and Patterson 2005; Smith and O’Brien 2005; Seldin 2007).

The high levels of LD in recently admixed populations are due to associations between pairs of loci co-inherited on an intact chromosomal chunk from one of the ancestral source populations(Chakraborty and Weiss 1988). This particular type of admixture-introduced LD, or ALD, decays as a function of time since admixture because of recombination. Consequently, it is possible to infer population admixture by modeling the dynamic changes of ALD. Patterson N. et al. recently proposed such an approach by aggregating pairwise LD measurements through a weighting scheme (Patterson *et al.* 2012).This was further developed by Loh et al.(Loh *et al.* 2013) This ALD based approach is particularly useful for admixture dating.

In the hybrid isolation (HI) model, the expected value of LD decreases at the rate of 1–*d* (Chakraborty and Weiss 1988; Pfaff *et al.* 2001), where *d* is the genetic distance (in Morgans) between two sites and *g* (generation) is the time elapsed since the initial admixture event. For convenience, (1–*d*)^*g*^ is considered approximately equal to *e*^−*dg*^ in computation, assuming that the admixed population is engaged in random mating and has infinite effective population size(Hill and Robertson 1966).

Pickrell et al. considered the situation of multiple waves of admixture from different source populations and showed that LD was comprised of multiple exponential terms, each of which refers to a single admixture event(Pickrell *et al.* 2014). However, the confounding effect was not fully taken into account, especially the effects of source populations on the LD of an admixed population. In this study, an accurate mathematical expression of LD is given under a general admixture model(Verdu and Rosenberg 2011) (Figure 1). Our analysis showed that the effect of the LD of source populations constitutes the major part of the LD in an admixed population (Table 1), which should not be ignored in the case of modeling or estimating ALD from empirical data.

**Table 1:** Absolute ratio of LD from source population to the LD of the admixed population

| Bin interval<br>(Morgan) | Raw LD |  | Average<sup>#</sup> Raw LD |  | Average Weighted LD |  |
| --- | --- | --- | --- | --- | --- | --- |
|  | Prop<sup>&</sup> | Median<sup>§</sup> | Prop | Median | Prop | Median |
| [0, 0.05) | 39.4% | 0.48 | 39.4% | 0.36 | 39.4% | 0.032 |
| [0.05, 0.1) | 12.4% | 0.45 | 12.4% | 0.33 | 12.4% | 0.028 |
| [0.1, 0.15) | 7.1% | 0.46 | 7.1% | 0.33 | 7.1% | 0.028 |
| [0.15, 0.2) | 6.7% | 0.45 | 6.7% | 0.33 | 6.7% | 0.028 |
| [0.2, 0.25) | 6.3% | 0.45 | 6.3% | 0.33 | 6.3% | 0.029 |
| [0.25, 0.3) | 6.1% | 0.45 | 6.1% | 0.33 | 6.1% | 0.029 |
| [0.3, 0.35) | 5.8% | 0.45 | 5.8% | 0.33 | 5.8% | 0.028 |
| [0.35, 0.4) | 5.6% | 0.45 | 5.6% | 0.33 | 5.6% | 0.028 |
| [0.4, 0.45) | 5.4% | 0.45 | 5.4% | 0.33 | 5.4% | 0.028 |
| [0.45, 0.5) | 5.3% | 0.45 | 5.3% | 0.33 | 5.3% | 0.027 |

For each value of pairwise LD, there is a corresponding genetic distance between the pair of SNPs. For convenience, the LD value and the corresponding genetic distance are called as the LD point. According to the value of the genetic distance, the LD point can be placed in bins.
<sup>&</sup>: Proportion of the number of LD points in the bin to all the number of LD points.
<sup>§</sup>: Median value of the ratio measured using Eq 17 with the LD points in each bin.
<sup>#</sup>: Average values are calculated from raw or weighted LD. First, the LD points were sorted according to the genetic distance. Then, with every 100 sorted LD points, the averages of the LD value and genetic distance were taken as the LD value and genetic distance as the new LD points. At last, these new LD points were placed in the bins and the proportion and median were calculated.

**Figure 1:**
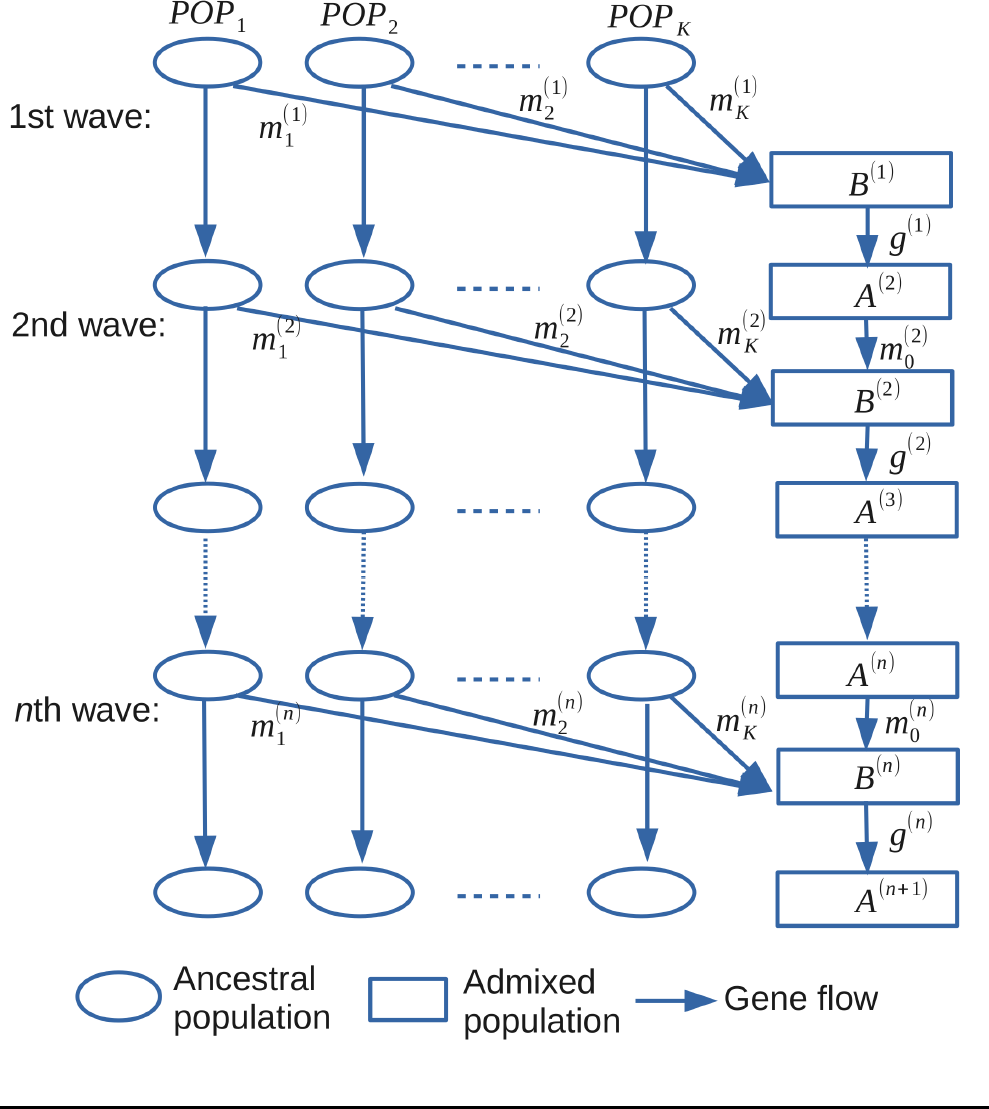
A general admixture model with K source populations and n waves of admixture. See Table S1 for notation.

Based on our mathematical description of the weighted LD, Eq 14, we proposed a new approach to separate the confounding effects of LD of source populations under a two-way admixture model. This approach was implemented in MALDmef, which was newly developed for this study. Unlike previous methods of estimating the time of admixture based on weighted LD, such as ALDER or MALDER, our method considers the influence of LD from source populations and there is no need to set the starting distance d_0_. The current method also involves a fast numerical method to fit the LD to hundreds of exponential functions to cover the admixture events across a range of hundreds of generations. Applying this method to the well-known admixed populations in HGDP(Rosenberg *et al.* 2002) and HapMap(Altshuler *et al.* 2010) data demonstrated that the current study have greatly facilitated understanding of the admixture history of human populations.

## Materials and Methods

### Data sets

Data for simulation and empirical analysis were obtained from two public resources: Human Genome Diversity Panel (HGDP)(Rosenberg *et al.* 2002) and the International HapMap Project phase III (Altshuler *et al.* 2010). Data filtering was performed within each population: Samples with missing rate > 5% per individual, SNPs with missing rate > 50% and SNPs failing in Hardy-Weinberg Equilibrium test (p-value < 1.0E-6) were permanently removed from subsequent analysis. Source populations for simulations used the haplotypes from 113 Utah residents with Northern and Western European ancestries from the CEPH collection (CEU) and 113 Africans in Yoruba (YRI).

### LD for general admixture model

A mathematical description of LD decay based on a general admixture model is given here (Figure 1). The relevant notation is summarized in Table S1. To produce the current model and render deduction more general and realistic, the population was assumed to be finite in size, and the effect of natural selection, if exists, is negligible. The details of math derivation can be found in Supplementary Material. However, the math expression of LD under the general assumption (Eq S10) is too complicated to be used in empirical analysis. It was here assumed that all populations were of infinite population size since the initial admixture. This assumption leads to a brief version of LD and it would have only limited effect on admixture time inference in empirical data analysis.

When the population size is infinite, the allele frequency on each site should always be considered as constant in the source populations: so for the *i*^*th*^ source population at the *l*^*th*^ wave admixture on the site *x* (Eq 14) and the allele frequency difference, Eq 14, between two source populations *i* and *j*:

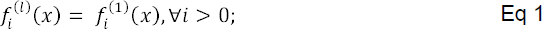

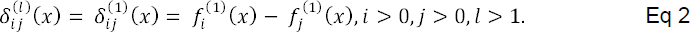

The total genetic contribution from source population *i* in the admixed population with *n*^*th*^ waves of admixture is as follows:

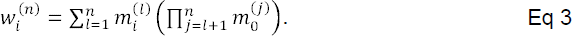

In this way, the expectation of LD for the target admixed population is as follows:

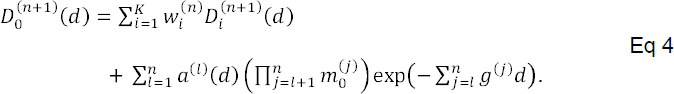

If we set *g*^*(j)*^ = 1 and

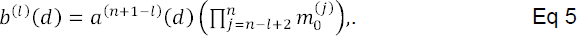

then we have the expected LD value for the *(n+1)*^th^ generation’s admixed population is as follows:

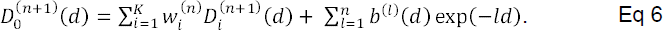

This equation tells us that the LD in admixed population is composed of two parts: one is from its source populations, and the other is formed by the admixture events.

### LD for specific admixture models

The mathematical description of LD for a general admixture model is provided above. Here, the LD of specific previously reported admixture models are described(Chakraborty and Weiss 1988; Ewens and Spielman 1995; Pfaff *et al.* 2001; Guo and Fung 2006; Jin *et al.* 2012). It is shown that these models can be regarded as special cases of our general model when parameters are specified (Tables S2-6).

The two-way hybrid-isolation (HI) model is the most popular admixture model and most admixture analyses are based on this model. Under this model, only two source populations are assumed to be involved in any one admixture event, and all the populations are isolated without further gene flow. In the current mathematical framework, description of LD for two-way HI model is as follows:

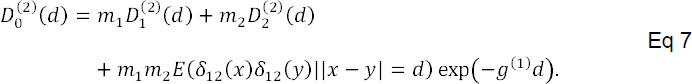

Two typical continuous admixture models have also been frequently discussed. These include the gradual admixture (GA) model(Ewens and Spielman 1995; Guo and Fung 2006) and continuous gene flow (CGF) model(Pfaff *et al.* 2001). Here, the two-way admixture of these models is discussed, but LD for the multiple-way admixture models are also incorporated into the current general admixture model. In these admixture models, we set the number of generations between two admixtures to be 1, *g*^*(j)*^ = 1 and the LD for the (*n*+1)^th^ generation of the admixed population (admixture began at the 1^**st**^ generation) is given below.

Under the GA model,

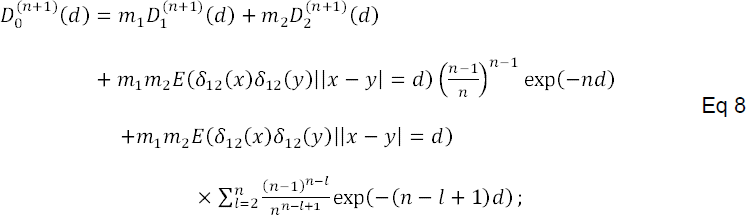

Under the CGF model,

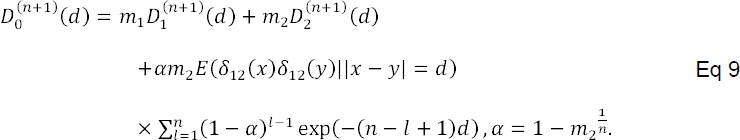

The GA model describes a scenario in which both source populations continuously contributed genetic materials to the admixed population after the admixture event, and the admixture proportion remained constant throughout all of the generations. However, the CGF model only allows one of the two source populations to continuously contribute genetic materials to the admixed population.

The two-wave admixture model describes that the admixed population is formed by two waves of admixture, one ancient and one recent. The ancient admixture event produces the admixed population and the recent admixture causes new migration in the admixed population. The LD (Eq S1) in the admixed population is as follows:

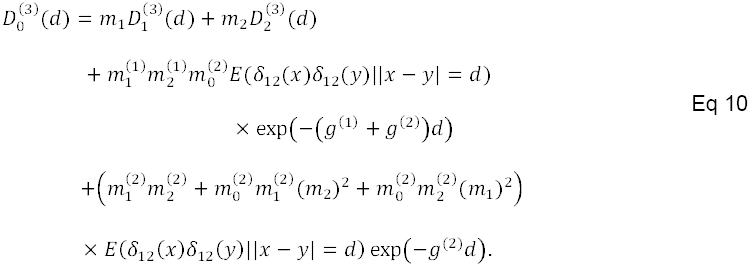

In a more complicated scenario, the genetic materials of the admixed population are inherited from more than two populations. One general scenario could be that two of the source populations (populations 1 and 2) meet first and then the third source population (population 3) joins the admixed population during the second wave. This phenomenon is here called the three-way-two-wave model. The LD in the admixed population under this model is as follows:

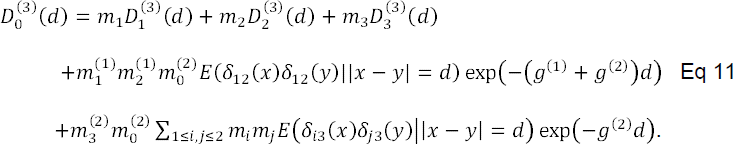

In order to provide an intuitive understanding of the dynamic decay of LD among these models, they were plotted with specific parameters (Figure S1).

### Modeling LD with consideration of influence of ancestral source populations

In order to demonstrate how the LD in admixed population is influenced by its ancestral source populations, we take the two-way HI model was here used as an example to illustrate the effect of the LD of source populations on the LD in the admixed population. Because, in a two-way HI model, no recombination occurs in the population immediately after admixture, the LD (Eq S2) in the first generation admixed population can be expressed as follows.

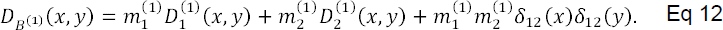

Here, the absolute ratio, *r*_sa_, to measure the effect of the LD in the source populations, is defined as follows:

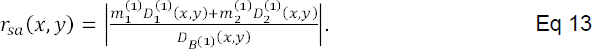

Here, the admixture between Europeans and Africans was considered. The scenario started with 100 individuals of CEU and 100 individuals of YRI as the source populations of Europeans and Africans. They were simply put together and treated as a population immediately after admixture. Pairwise LD was calculated using the sampled SNPs on chromosome 1. The SNPs were sampled in two rounds. In the first round, one SNP was sampled for every twenty SNPs. In the second round, SNPs on the right side of every SNP sampled in the first round (with greater physical position numbers) as follows: All the SNPs whose genetics distance from the chosen SNP were below 0.05 Morgans were sampled; one in every three SNPs in the region whose genetics distance from the chosen SNP was between 0.05 and 0.1 Morgans were samples; and one in every five SNPs in the region whose genetic distance from the chosen SNP was between 0.1 and 0.5 Morgans were sampled. LD was calculated only between the SNPs sampled in the first round and the related SNPs sampled in the second round. The *r*_sa_ was calculated with raw LD of YRI, CEU and the manufactured admixed population and these were classified into bins according to the genetic distance between each pair of SNPs. Results showed the median value of *r*_sa_ to be around 0.45 for all bin intervals (Table 1). The LD of source populations made up about 45% of the LD in source populations. However, recent work with the LD provide insight into population admixture history, even though the effect of LD from the source populations was ignored(Loh *et al.* 2013).

Actually, using the average value of weighted LD can reduce the effect from source populations. To verify this conclusion, we first calculated *r*_*sa*_ on the average value of LD (for every 100 pairs of SNPs) in each bin (classified according to the genetic distance between each pair of SNPs) and then calculated *r*_sa_ on the average value of weighted LD in each bin. Results showed LD from the source populations to make up about 33% of the average LD and only about 2.8% of the average weighted LD in the admixed population, and this proportion did not deduce to be negligible as the genetic distance increasing (Table 1). In summary, this analysis indicates that weighted LD is more proper to be used to infer admixture because it contains less effect of confounding LD. However, using a starting distance with weighted LD might not be the best way to reduce the confounding effect of source populations.

### Weighted LD in a two-way admixed population

The effective population size of the admixed population was assumed to be large enough to be regarded as infinity, the LD can be described by Eq 6. The effect of admixture events could be formalized with the sum of bunches of exponential functions. The coefficient of exponential functions, Eq 6, if positive, indicates the admixture event l generations ago. This can be used to estimate the time of admixture. Moreover, it gives us an opportunity to infer the history of admixture without any prior assumption of particular admixture models (such as HI, GA, or CGF). In this way, it is important to study the exponential property of the LD decay in admixed populations. If the ancestral source populations are known for the two-way admixed population, the LD is as follows:

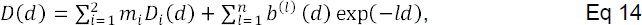

Here, *D(d)* is the LD of the admixed population, *m*_*i*_ is the genetic contribution from source population *i* to the admixed population and *D*_*i*_*(d)* is the present LD of the ancestral source population *i*.

To determine the time of admixture actually using the exponential term *exp*(−*ld*), two important things must be taken into consideration. First, the coefficient of the exponential function, *b*^(*l*)^(*d*), must be relatively constant and bigger than 0; second, the effect of the LD of source populations can be separated. This confirmed that using weight, and the difference in allele frequencies between two source populations, could leave the coefficients of the exponential functions constantly bigger than zero (Appendix) and it may be possible to deduce the effect from source populations (Table 1). The weighted LD is examines in further detail below. The formula of weighted LD is as follows:

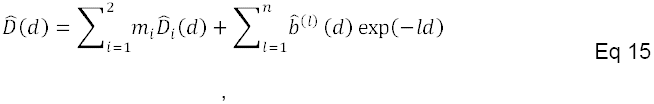

Here,

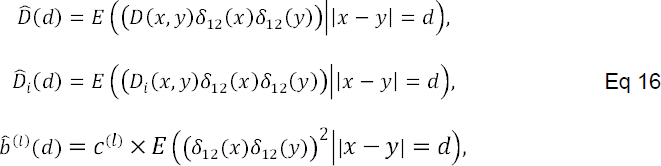

and

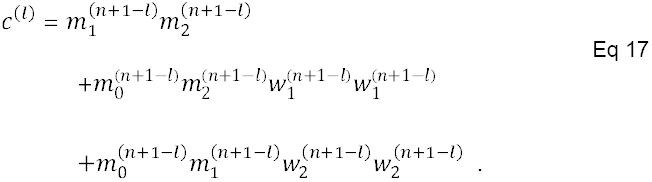

Here, *c*^(*l*)^ is determined by the admixture history and it is a natural indicator for admixture events.

### Factorizing of weighted LD with exponential functions

In order to show the exponential properties of the weighted LD, the admixed population was simulated using forward time simulation with haplotype data of YRI and CEU. A 100-generation old admixed population with 50%:50% proportion was constructed. Ancestral source populations based on the haplotype data of YRI and CEU were also constructed in the simulation, separately. Next, 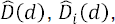, and 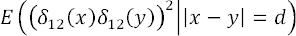 were calculated based on the genotype data. The LD decay was fit with hundreds of exponential functions. In this way, the coefficient spectrum on exponential functions was determined and used to describe the decay of weighted LD. The fitting method was able to model the decay of weighted LD well and to provide the amplitude in every exponential function. The results of fitting 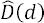 primarily showed three bunches of exponential functions composing the LD decay, with *l* values around 100, 180, and 1,250 generations (Figure 2A). As mentioned above, the *l* value corresponds directly to the time of admixture, so the fact that the signal was around 100 could be explained easily by the admixture in that the designed admixture is 100 generations ago.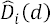 and 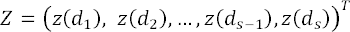 were also fit, and both 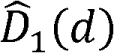 and 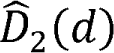 showed signals on *l* around the value 1,250 (Figures S2–3). No significant signal peak was observed with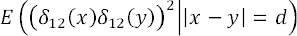, even though there was a sharp decay over a very short distance (Figure S4). Based on these fitting results, it is here speculated that the signals in 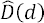 around 1,250 and 180 might have resulted from confounding LD of the ancestral populations. To test this hypothesis, *Z*(*d*), the optimized ALD, was fit so that LD of source populations was separated from the admixed population as the follows:

**Figure 2:**
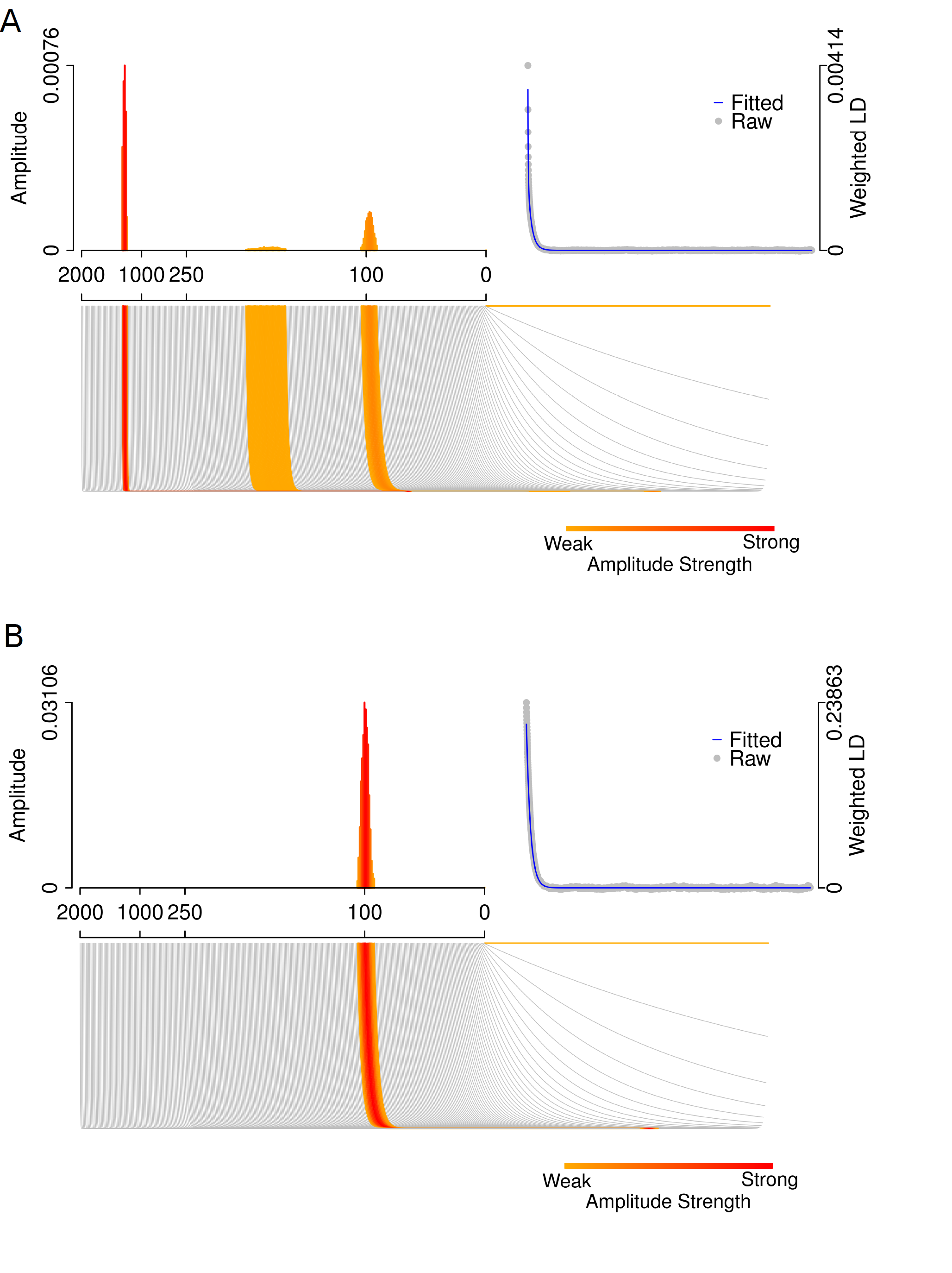
Full exponential spectrum for fitting extending weighted LD in a simulated admixed population. A) Exponential spectrum without separating the confounding LD from the source populations. B) Exponential spectrum with separating the confounding LD from the source populations.

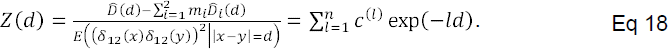

The fitting results of *Z(d)* showed only signal peak around the 100 of *l*’svalue (Figure 2B). This supported the present hypothesis.

In summary, the weighted LD of source populations may affect the exponential properties of weighted LD of admixed population for relatively large *l* values, which may be the cause of signals of ancient admixture when employing a LD-based method for time estimation. However, we can use the derived LD of source populations to reduce the fake admixture signals.

### Calculation of weighted LD and fitting LD decay

The basic algorithm used to calculate weighted LD was the same as that used with ALDER and MALDER. It was coded in C++. The weighted LD for admixed population was calculated as follows:

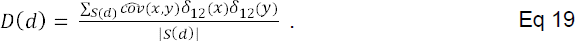

Here,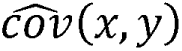 is a non-bias estimator of the covariance on the genotype between two SNPs at *x* and *y*. S(d) is the set holding pairs of SNPs with inter-site distance of d Morgan. *δ*_12_(*) is the allele frequency difference, defined in Eq S3. The weighted LD for the source populations are calculated as follows:

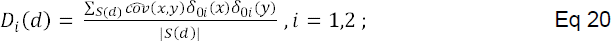

Here, ‘0’ represents admixed population.

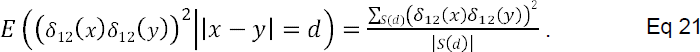

In this way, the algorithm and calculations relied only on genotype data, which prevented the introduction of phasing errors. The fast Fourier transform algorithm was also used to increase the computational efficiency. The population admixture proportions were estimated using the following formula.

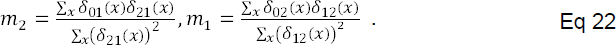

Once it is possible to calculate *Z(d)*, we can fit it using a numerical routine known as the proximal gradient(Beck A. amd Teboulle *et al.* 2009). The object function to minimize is as follows:

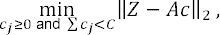

where Z = (*z*(*d*_1_), *z*(*d*_2_),…, *z*(*d*_*s*−1_),*z*(*d*_*s*_))^*T*^ is a vector of *Z(d)* with different d values. These can be obtained from genetic data. *c* = (*c*_0_,*c*_2_,…,*c*_*n*−1_, *c*_*n*_)^*T*^ is the coefficient of the exponential functions. The *(i,j)^th^* entry of the matrix *A*_*s×(n+1)*_ is *A*_*ij*_ = exp(*−d*_*i*_*G*_*j*_), where {*G*_*j*_}_*j* = 1,..,*n*+1;_ is the chosen subset of data in the generations from 0 to *G*. With this method, it was possible to find the positive values in the vector *c*.

### Determination of the significance of admixture signals and denoising

The significance of admixture signal *c* was measured using a Jackknife-based approach. For the target population, each chromosome was excluded one at a time and the value of *Z(d)* was calculated using the remaining chromosomes. After fitting *Z*(*d*) with the sum of exponential functions, denoising was performed on the coefficients of exponential functions. Only the top signals that composed 99.9% of *Z*(*d*) were retained. For the coefficient of each exponential function, 22 observed values were tested to determine whether they were larger than 0. In this way, P-values were used to measure the significance of the admixture signal. In the spectrum plot of coefficients, the mean of the 22 coefficients with P-values smaller than 0.05 were plotted for each exponential function. This was specified using the candidate admixture time points. (Figure 2 and Figure 5)

### Simulations

In order to evaluate the performance of our method for estimating the time of admixture, forward-time simulation was used to generate haplotypes of admixed populations under different admixture models and different scenarios: HI model, two-wave model (including the situation of one source population and the situation of two source populations which contribute genetic materials to the admixed population in the second wave), and continuous gene flow model were used in a combination of GA and isolation models and of CGF and isolation models. In our simulation, the newly generated haplotypes are assembled with the segments of haplotypes in source populations(Li and Stephens 2003; Price *et al.* 2009), which in this particular case come from the haplotypes of CEU, and YRI.

In the HI model, the admixture event was set as having occurred 100 generations earlier. In the two-wave model, the first admixture event was set as having occurred 100 generations earlier, after which the admixed population was isolated for the next 80 generations, until the second wave of admixture, after which the admixed population was isolated another 20 generations. In the second admixture event, a scenario in which only one of the source populations donated genetic materials (TW-CGF model) and another scenario where both source populations provided gene flows (TW-GA model) were simulated.

Scenarios with continuous migration were also simulated. In one scenario, the migration window was set to 80 generations: Continuous gene flow was simulated from one of the source populations was simulated for 80 generations and then isolated for the next 20 generations (CGF-I model), and continuous gene flow was simulated from both of the two-source populations for 80 generations and then the admixed population was isolated for 20 generations (GA-I model); in the other scenario, the migration window lasted 30 generations: 30 generations’ continuous gene flow and 70 generations’ isolation were used in both the CGF-I model and GA-I model. The evolution of the source populations was also simulated under random mating with sample size of 5,000 for 100 generations. The parameter details are given in Table S7–8.

## Results

### Robustness of the new method when used on proxy source populations

A way of estimating the time of admixture was developed by separating the LD from source populations directly from that of the admixed population, which requested us to know the true ancestral source populations. However, identifying the true source populations is also a complicated and difficult problem. In most cases, only limited populations are available. In the appendix, it is shown that using the proxy populations similar to the true source populations can show the exponential properties of weighted LD’s decay well. It is here claimed that using proxy populations can also reduce the confounding effect attributable to source populations’ LD. Here, YRI and CEU served as source populations from Africa and Europe, respectively, and 100 admixed individuals were simulated using 100 generations’ admixture. This method was able to show the admixture time with the true source populations and pairs of source populations representing different parts of Africa and Europe, i.e. CEU-LWK, CEU-MKK, TSI-LWK, TSI-MKK, and TSI-YRI, very well (Figure S5–9).

### Robustness of the new method in various admixture models

This method also involved using different admixture models. In order to render the results of the evaluation reliable, 10 independent admixed populations with haplotypes obtained from 113 unrelated CEU individuals and 113 unrelated YRI individuals were simulated with data from all of 22 autosomes. When the weighted LD was calculated, 100 individuals were sampled from current source populations (isolated for 100 generations with random mating) and admixed population separately. Under various admixture models, MALDmef was able to reconstruct the history of the admixture population well. For the one-pulse and two-pulse admixture models, MALDmef gave the time close to the true time of admixture for the continuous migration models, MALDmef was able to place most of the signals in a particular migration time interval (Figure 3).

**Figure 3:**
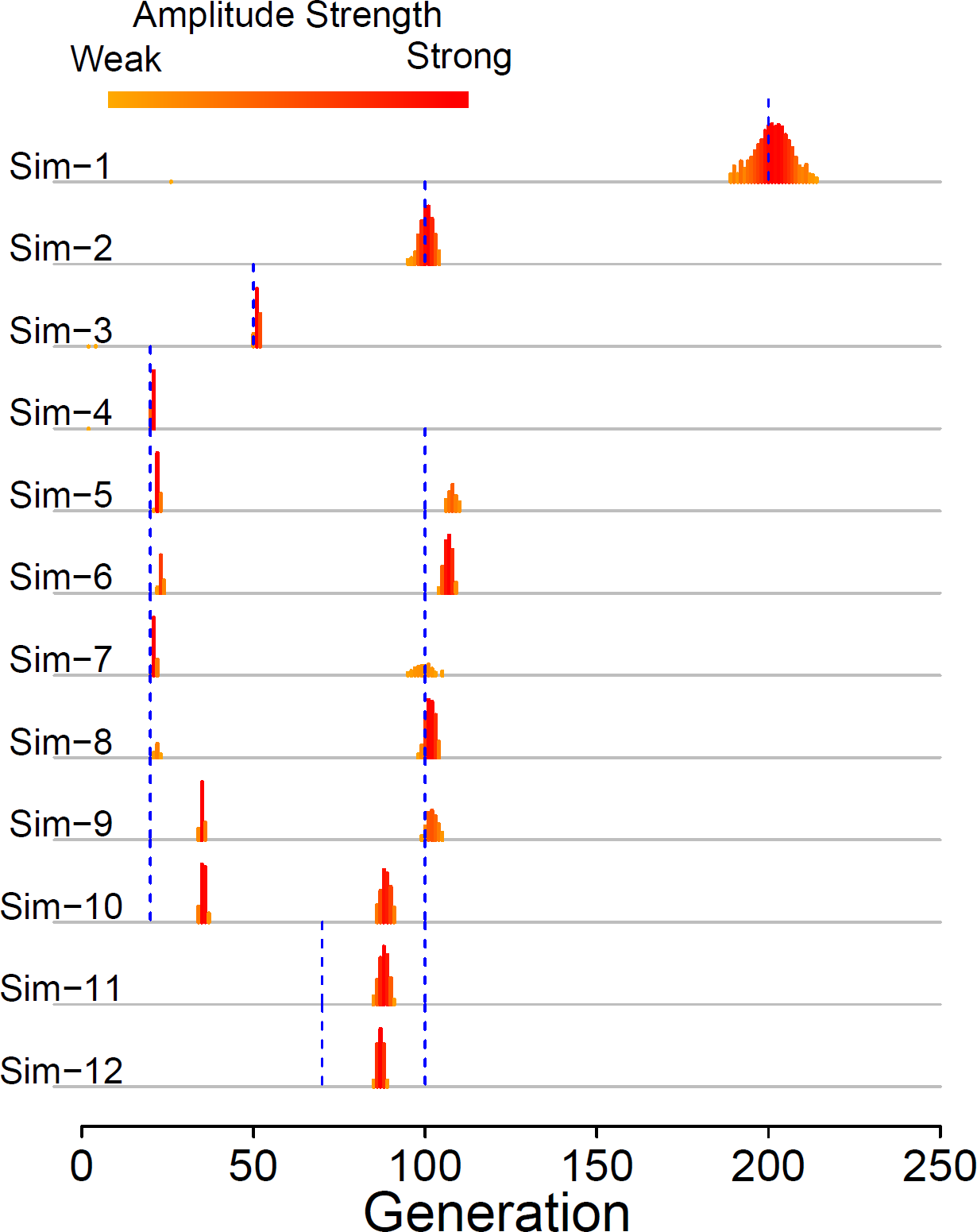
Evaluation of the performance of MALDmef under various admixture models. The blue vertical dash line represents the true simulated admixture time. Sim-1 to Sim-4: one-pulse admixture of 200, 100, 50 and 20 generations ago. Sim-5 to Sim-8: two-pulse admixture at 20 and 100 generations ago. Sim-9 to Sim-12: continuous admixture from 20 to 100 generations ago (Sim-9 and Sim-10) and from 70 to 100 generations ago (Sim-11 and Sim-12).

MALDER was also run on the same simulation data. MALDER is the only software that can deal with multiple-wave admixture. CHB and CHD were selected as the extra source populations and the starting distance was set to 0.005 Morgans and the bin size was set to 0.0002 Morgans. Under the HI model, our method revealed significant signals around the generation (100) we set for simulation (Figure S10), while MALDER gave us 8 significant signals of 10 independent simulations around 100 generations. Here, 5 of the 8 signals were accompanied by significant signals above 250 generations, which could be caused by the LD from the source populations (Figure S11). For the multiple-wave admixture, we defined an estimation deviation (ED) to measure difference (distance) of the location between the true admixture signals and the detected admixture signals, which is defined as follows:

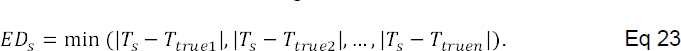

Where {*T*_*truej*_,*j* = 1, 2,3,… *n*} is the set of true admixture time and {*T*_*s*_} is the estimated times of admixture. In our simulations, the following was true:

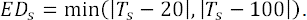

The results under HI model can also be estimated as follows

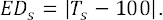

Under repeating simulations for the other various admixture models, the *ED* values of MALDmef were significantly smaller than the *ED* values based on MALDER, indicating that the current method is more precise and stable than MALDER (Figure 4). The details of estimation with MALDER and MALDmef on repeating simulations are shown in Figure S10–27. The current method was also used on the empirical admixed populations.

**Figure 4:**
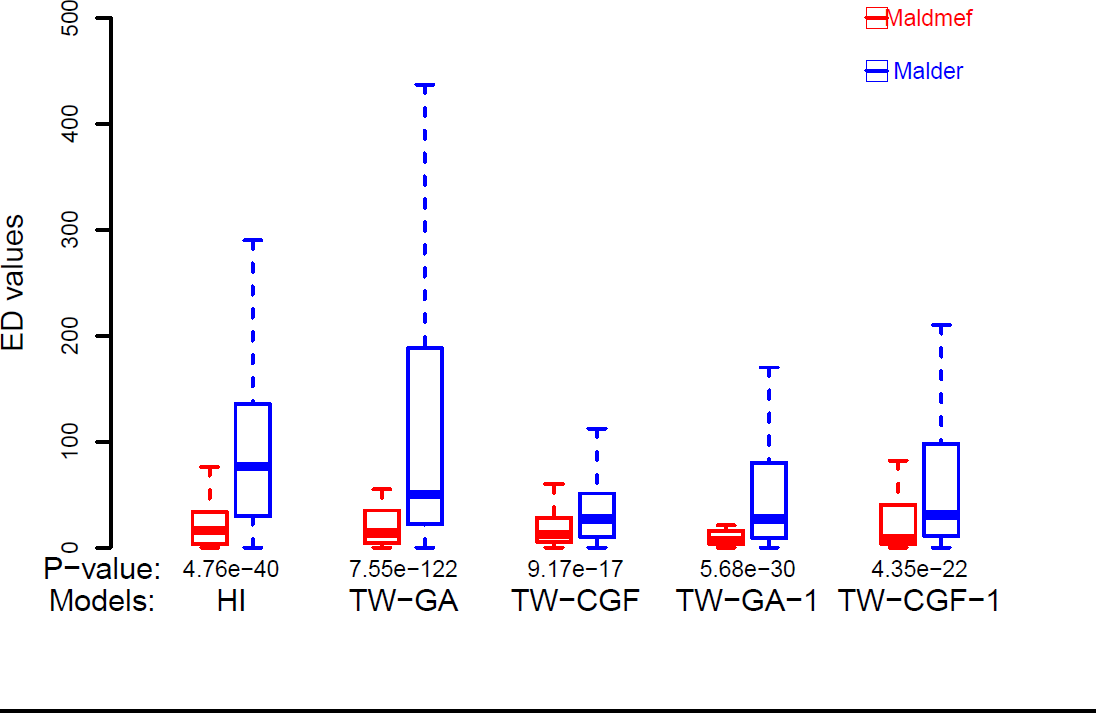
Performance of MALDmef and MALDER in various simulations. One-side t-test was performed to calculate the significance of the differences between ED values of MALDmef and MALDER. The detailed parameters for each simulation are given in Table S7.

**Figure 5:**
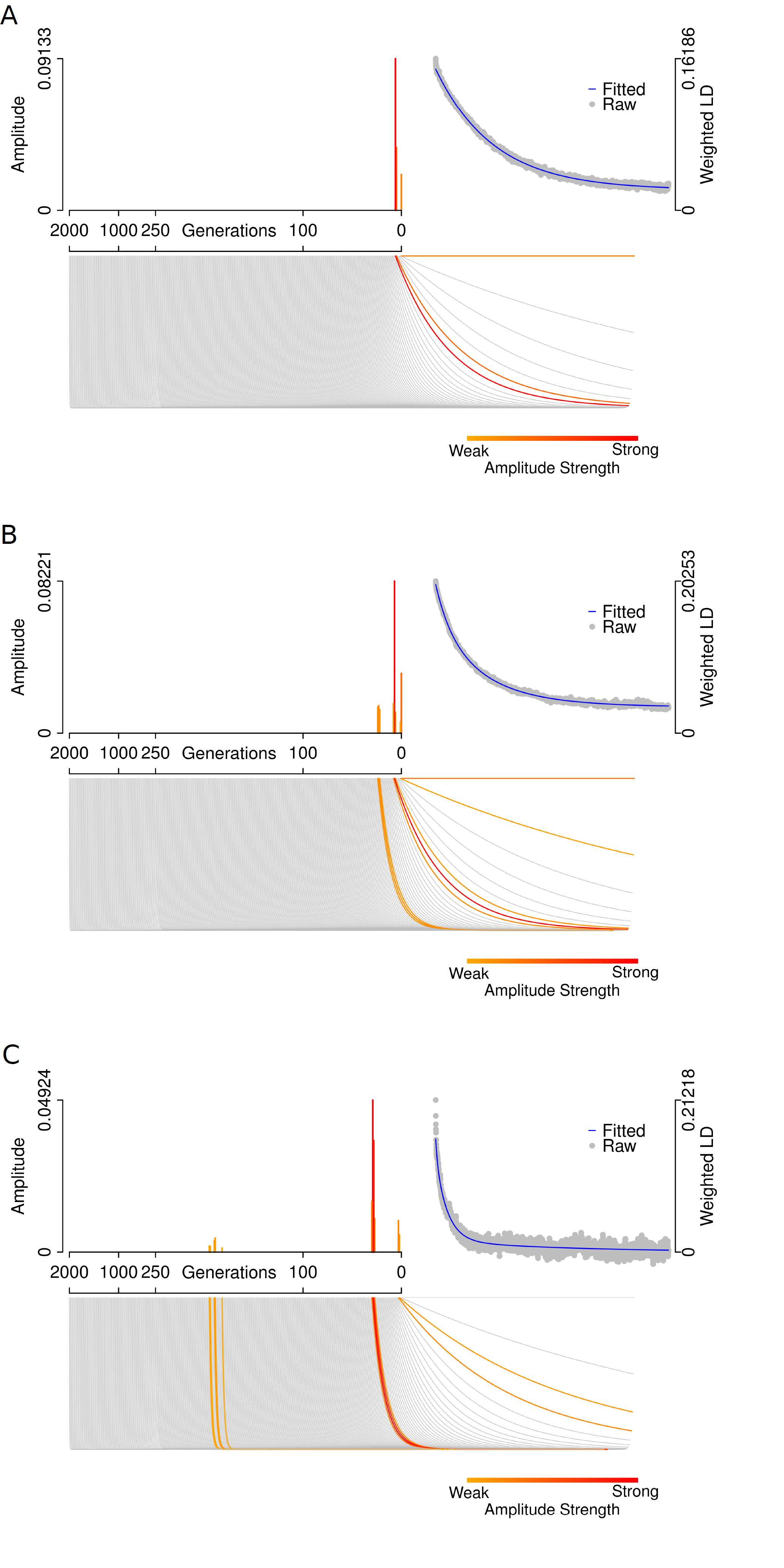
Full exponential spectrum for fitting extending weighted LD in admixed populations. A) ASW; B) MEX; C) Uyghur.

### Estimating admixture time using empirical data

The current method was first applied to a few well-known admixed populations as in analysis reported by Alder(Loh *et al.* 2013) using an available public database(Rosenberg *et al.* 2002; Altshuler *et al.* 2010). MALDmef can currently only deal with the two-way admixture when derived source populations or with the populations similar to the true derived source populations are available. However, the real admixture history could be much more complicated than assumed and many factors may affect the results of estimation. In order to interpret the time spectrum, three principles should be followed:

1. A signal with larger amplitude is more reliable than one with smaller amplitude.
2. A signal that remains on the time spectrum for longer than 250 generations indicates that the chosen source populations are probably not similar enough to the real ancestral source populations or that the general two-way admixture model does not fit the data well.
3. A signal in the continuous time interval containing generation 1 may reflect the substructure of the admixed population but not the admixture.

Based on these principles, MALDmef was first applied to the well-known admixed populations: African American (57 ASW individuals from HapMap), Mexican (86 MEX individuals from HapMap) and Uygur (10 Uygur individuals from HGDP). MALDER was also used to analyze these admixed populations.

In our analysis with MALDmef, CEU (n = 113) and YRI (n = 113) were chosen as the ancestral populations of ASW. CEU (64 individuals) and American Indian (7 Colombians, 14 Karitiana, 21 Maya, 14 Pimas and 8 Suruis) were chosen as the ancestral populations of MEX. Han (n = 34) and French (n = 28) were chosen as the ancestral populations of Uygur. The time of admixture of ASW was found to be about 4 to 5 generations ago (100–125 years before present, assuming 25 years per generation) (Figure 5A). MEX seems to experience two wave of admixture: ranging from 6 to 8 generations (150–200 years) ago and from 22 to 24 generations (550–600 years) ago, respectively (Figure 5B). Neither inferences on ASW nor those on MEX showed any significant signals on the time earlier than 250 generations ago, suggesting that the source populations chosen for ASW and MEX are similar enough to the true derived ancestral populations. The Uygur population has been reported to have much longer admixture history than ASW and MEX(Xu and Jin 2008; Xu *et al.* 2008; Jin *et al.* 2012; Qin *et al.* 2015). It showed three time intervals of admixture: 2 to 3, 27 to 30, and 182 to 195 (182,189,190,194,195) (generations before present) (Figure 5C). The most significant signals lay in the interval of 27–30, suggesting that the major admixture creating the current population happened around 575 to 750 years ago, which is consistent with previous result on the recent admixture. However, the signals on the time interval from 182 to 195 generations ago may indicate the ancient admixture events. Because of the sample size limitation of LD-based method, an accurate inference on the ancient wave of admixture may be difficult to find for Uygur. Private screening of 92 fully sequenced individuals of Uygur indicated two highly confident admixture intervals: from 18 to 20 generations ago and from 83 to 95 generations ago, supporting the conclusion that an ancient admixture event helped create the modern Uygur.

Loh et al. speculated that there could have been multiple waves of admixture in the history of MKK (Loh *et al.* 2013). In the current analysis, 113 YRI and 113 CEU individuals were used to represent the source populations of the MKK (n = 156). Here 5 candidate intervals of admixture were identified, having occurred: 13–16, 50–67, 107–139, 310–410, and 1190–1210 (generations before the present). Only three of them, 13–16, 50–67, and 107– 139, seemed to be admixture signals, indicating that the three admixture events happened from 325–400, 1,250–1,675 and 2,675–2,475 years ago. The signals from between 1190 and 1210 generations ago may also indicate that the ancestral populations are not good enough to infer the time of admixture (Figure S28)

Paralyzed analysis with MALDER was also conducted on these admixed populations (Table 2). For each admixed population, first all the populations in the full data were set as the reference populations to infer the admixture, and then the pair of populations with the highest amplitude for each wave of admixture were collected and used as the reference populations to re-run MALDER. MALDER showed good consistency in two rounds of inference. From the results with global populations as references, MALDER was able to determine the pair of populations that fit each wave of admixture best. Then we MALDmef was run with each pair of populations as the reference to the corresponding admixed population (Table3). Results showed that MALDmef was also very robust to the reference populations and it showed considerable consistency in each wave of admixture. However, with different reference populations suggested by MALDER, it detected signals from even earlier than 250 generations back. This result indicated that those populations suggested by MALDER might not be the proper source populations to study the admixture under a two-way model.

**Table 2.** Time of admixture (generations) determined using MALDmef with selected reference populations.

| Admixed population | Reference populations | Time 1 | Time 2 | Time 3 | Time 4 | Time 5 |
| --- | --- | --- | --- | --- | --- | --- |
| ASW | CEU;YRI | 4-5<br>$2.02 \times 10^{-13}$ | | | | |
| MEX | CEU;American Indian | 6-8<br>0.00506 | 22-24<br>0.00561 |  |  |  |
| MEX | American Indian; YRI | 8-9<br>$3.38 \times 10^{-5}$ | 32-38<br>0.0481 | 810-830<br>0.00129 | | |
| MKK | CEU;YRI | 13-16<br>0.0205 | 50-67<br>0.0456 | 107-139<br>0.0499 | 310-410<br>0.0284 |  |
| MKK | TSI;YRI | 17-19<br>0.00271 | 64-79<br>0.0337 | 115-160<br>0.0494 | 185-188<br>0.0481 | 590-670<br>0.0147 |
| Uyghur | French;Han | 2-3<br>$1.82 \times 10^{-6}$ | 27-30<br>0.0448 | 182-195<br>0.0484 | | |
| Uyghur | Basque;Han | 2-4<br>0.0385 | 29-31<br>0.00377 | 157-164<br>0.0427 | 690-750<br>0.0441 |  |
| Uyghur | Sardinian; Japanese | 2-3<br>$2.79 \times 10^{-6}$ | 32-35<br>0.0149 | 209-211<br>0.0421 | 710-790<br>0.0253 | |

3 In each time cell, the biggest P-value is provided under the detected time 4 interval.

**Table 3:** Time of admixture (generations) determined using MALDER with global scanning and selected populations

| Admixed pop | Reference pop | Time 1 | Time 2 | Time 3 | Time 4 |
| --- | --- | --- | --- | --- | --- |
| ASW | HapMap | 2.7 +/- 0.5<br>(Z=5.27)<br>CEU;YRI | 12.0 +/- 4.4<br>(Z=2.76)<br>CEU;YRI |  |  |
| ASW | CEU;TSI;YRI | 1.3 +/- 2.8<br>(Z=0.47)<br>CEU;YRI | 6.1 +/- 3.2<br>(Z=1.92)<br>CEU;YRI | 70.3 +/- 60.7<br>(Z=1.16)<br>TSI;YRI |  |
| MEX | HapMap+<br>American<br>Indian (HGDP) | 5.2 +/- 1.9<br>(Z=2.78)<br>Indian;TSI | 19.8 +/- 9.7<br>(Z=2.05)<br>Indian;YRI |  |  |
| MEX | American<br>Indian; TSI;YRI | 4.2 +/- 1.6<br>(Z=2.67)<br>Indian;TSI | 15.1 +/- 4.0<br>(Z=3.82)<br>Indian;YRI |  |  |
| MKK | HapMap | 3.3 +/- 0.7<br>(Z=4.51)<br>TSI;YRI | 23.7 +/- 6.4<br>(Z=3.69)<br>CEU;YRI | 107.8 +/-<br>32.7 (Z=3.29)<br>TSI;YRI | 495.2 +/- 255.1<br>(Z=1.94)<br>TSI;YRI |
| MKK | YRI;CEU;TSI | 3.1 +/- 0.8<br>(Z=3.87)<br>TSI;YRI | 20.3 +/- 6.4<br>(Z=3.19)<br>YRI;CEU | 80.3 +/- 28.3<br>(Z=2.84)<br>TSI;YRI | 292.7 +/- 121.5<br>(Z=2.41)<br>TSI;YRI |
| Uyghur | HGDP | 4.3 +/- 2.4<br>(Z=1.79)<br>Basque; Han | 40.9 +/- 7.4<br>(Z=5.52)<br>Japanese;<br>Sardinian |  |  |
| Uyghur | Basque;Han;<br>Japanese;<br>Sardinian | 4.6 +/- 2.6<br>(Z=2.12)<br>Basque;Han | 41.4 +/- 7.0<br>(Z=5.92)<br>Japanese;<br>Sardinian |  |  |

3 For each wave of admixture, we listed the pair of populations with the biggest 4 amplitude value under the Z score.

The MALDmef’s results were comparable to MALDER’s on the same admixed populations. In recently admixed populations, such as ASW and MEX, MALDmef and MALDER had similar results but MALDmef had shorter time interval ranges. In the admixed population with related ancient admixture, such as Uygur, MALDmef was more powerful and stable than MALDER in detecting the ancient admixture. MALDmef was able to predict the possible ancient admixture events that produced the modern Uygur population. With MALDmef, this can be supported by large sample sizes and dense markers. This is not the case with MALDER.

## Discussion

A general admixture model was here used to demonstrate the dynamics of LD in admixed populations. A method was developed based on this model and applied to estimating the time of admixture of populations that has undergone two-way admixture, and for the first time, the admixture time spectrum was used to reveal numbers and times of admixture events having occurred in a particular population. This method and results should shed new light into the drawing of inferences in the population genetics of admixed populations.

In this study, results confirmed that the extent of LD was composed of multiple exponential curves as a function of genetic distance in a certain admixed population formed by multiple waves of admixture from multiple source populations. Moreover, the confounding effects of source populations were demonstrated using mathematical description of the LD.

In previous studies, LD from source populations was usually assumed to be negligible in admixture time inference(Patterson *et al.* 2012; Loh *et al.* 2013; Pickrell *et al.* 2014). However, it was here shown to make up a major part of the LD in admixed population, but the composition in the average values (in bins, determined by genetic distance) of weighted LD was rather small. The results of the current study showed the average proportion to be about 33% in the values of pairwise LD and the proportion to be about 2.8% in the average values of weighted LD. However, both these values decreased slightly as genetic distance increased. This indicated that using weighted LD could really help reduce the confounding effect of source populations, but choosing a starting distance might not be the best way to reduce the confounding effect of source populations.

Based on these considerations, the new method we developed in this study was able to infer the time for multiple-wave admixture. In this method, the reference populations were first used to reduce the confounding effect of true. This method was very robust when the true ancestral source populations were not available, and it worked well by using reference populations similar to the source populations. The admixture-induced weighted LD extent curve was fit with hundreds of exponential functions, which provided a signal spectrum on time points ranging from 0–2,000 generations, which were within the range of most possible admixture events after the migration of modern humans “Out of Africa”. The jackknife method was used to produce a *P*-value on each time point and then determined the time intervals for the possible admixtures. Using various simulations, this method was demonstrated to be more accurate in estimating admixture time especially in the scenario of two-way admixture than other available methods.

This method, MALDmef, was used to analyze simulated data generated with a continuous admixture model. Results could also be interpreted as multiple-wave admixture instead of the continuous admixture model, here simulated as the true model. This constitutes potential bias that could affect the interpretation of the results. Considering this issue, we propose a particular concept, namely **effective admixture**, i.e., continuous admixture that can be treated as a few single-pulse hybrid events. This treatment should at least make sense in the admixture modeling. The rationale is that the inference of admixture should be still valid even with a model assuming a scenario of hybrid-isolation, in a sense of critical parameters could still be estimated effectively, but the admixture process could not be assessed precisely. The method developed in this study, i.e., MALDmef, can be used to infer the effective admixture events.

In the current analysis, it was observed that the weighted LD (one reference) of source populations (no admixture events) could also fit exponential functions closely with the major signals indicating very ancient events, such as, 1,250 generations ago, but the mechanism has yet to be well explained.

Even though the current method infers the time of multiple-wave admixture under the two-way model, it still has many limitations regarding the complicated demographic history of admixed populations. For example, during time estimation with weighted LD, the effects of nature selection and inbreeding were not considered here; neither was the situation of multiple-way admixture. This was because the means by which the weight terms affect the LD in multiple-way admixture have yet to be worked out. However, the improved method developed in this study may facilitate the inference of admixture history, particularly in determining multiple-wave population admixture. Future work should focus on developing methods capable of inferring multiple-way admixture.

## Appendix

This appendix describes the properties of the coefficients of the exponential terms with or without difference in allele frequency as weights. The source populations are assumed to be infinite in size and to remain so after the first admixture, so the unweighted coefficient is as follows:

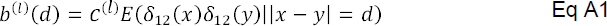

and the weighted coefficient is as follows:

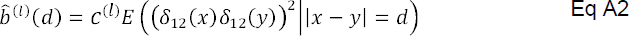

*c*^(*l*)^ is defined using Eq 17. Strictly speaking, both *b*^(*l*)^(*d*) 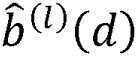are the functions of *d*, and they consist of two parts: The part in the first parenthesis is determined by the admixture; the other is the conditional expectation, determined by the allele frequency difference between the two source populations. The part determined by the admixture is independent of the genetic distance between two markers. It can be regarded as constant during the extraction of time information from the exponential terms. The part that matters is the conditional expectation. If the genetic distance *d* is large enough that *δ*_12_(*x*) is independent of *δ*_12_(*y*), the value of *E*(*δ*_12_(*x*)*δ*_12_(*y*)||*x* − *y*| = *d*) will be zero, but the value of *E ((δ_12_(x)δ_12_(y))^2^|x − y| = d)* should be a certain constant that measures the genetic distance between source populations 1 and 2. In this way, 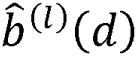 can help infer the exponential terms related to the admixture more effectively than *b*^(*l*)^(*d*)

Another problem addressed here is the property of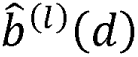, when only get the populations close to the true source populations were available to calculate the weighted LD. Suppose population 3 is similar to population 1 and population 4 to population 2, they follow a genealogical relationship described in Figure S29. The weight used here is as follows:

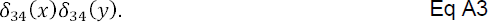

Then

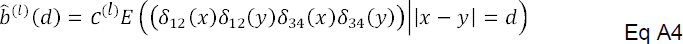

Under the same assumption, the genetic distance *d* is considered large enough that *δ*_12_(*x*) and *δ*_34_(*x*) are independent of *δ*_12_(*y*) and *δ*_34_(*y*).

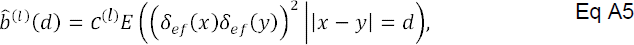

Here, *e* represents the shared ancestral population of 1 and 3 and *f* is the shared ancestral population of 2 and 4. In situation in which the true source populations are not available or cannot be identified, populations similar to the source populations can be used to calculate weighted LD.

### Software

C++ Source codes of MALDmef can be found at: http://www.picb.ac.cn/PGG/resource.php

## Acknowledgements

These studies were supported by the Strategic Priority Research Program of the Chinese Academy of Sciences (CAS) (XDB13040100), by the National Natural Science Foundation of China (NSFC) grants (91331204 and 31171218). S.X. is Max-Planck Independent Research Group Leader and member of CAS Youth Innovation Promotion Association. S.X. also gratefully acknowledges the support of the National Program for Top-notch Young Innovative Talents of The “*Wanren Jihua*” Project. The funders had no role in study design, data collection and analysis, decision to publish, or preparation of the manuscript.

